# AutoSpectral improves spectral flow cytometry accuracy through optimised spectral unmixing and autofluorescence-matching at the cellular level

**DOI:** 10.1101/2025.10.27.684855

**Authors:** Oliver T. Burton, Leoni Bücken, Leana De Vuyst, Stephanie Humblet-Baron, Anjana Lopez Menoz De Leon, Sameen Khan, Joana Cerveira, James Dooley, Adrian Liston

## Abstract

The advent of spectral flow cytometry has seen a rapid rise in the complexity of flow cytometry experiments, allowing the construction of assays with at least 50 fluorescent parameters. To correctly determine the contributions of each fluorophore’s signal to the high parameter data an accurate unmixing matrix needs to be generated. Even with single-stained controls, however, these matrixes include errors such as spillover spread, which compounds with each additional parameter, functionally limiting panel design. An additional source of errors is heterogeneity of cellular autofluorescence, which can affect both the unmixing matrix and misalign signals when the matrix is applied to individual cells in complex cell mixtures. Here we developed AutoSpectral, a statistical approach to automate the production of minimal-residual error unmixing matrixes and pair multiple distinct multifluorescent spectra to individual cells within a mixed sample, via an R-based software tool. AutoSpectral improves unmixing accuracy, improving incorrectly assigned cell positions by up to 9000-fold, reduces spread, particularly in samples with variable autofluorescence, and allows multi-lineage analysis of mixed populations, providing superior data for spectral flow cytometry experiments.

## INTRODUCTION

Spectral flow cytometry, with the capacity to unmix the signals from similar fluorophores, is a recent advance in the flow cytometry field. Spectral flow cytometry has unlocked a rapid increase in the number of parameters that can be simultaneously measured, dramatically increasing the information richness of datasets. Spectral unmixing, the deconvolution of aggregate fluorescent signals into individual fluorophore abundances reporting on marker expression, requires careful preparation of samples stained with examples of each fluorophore in isolation, called single-stained controls. Unfortunately, the data processing of these single-stained controls and generation of an optimised unmixing matrix have not kept pace with the hardware improvements of spectral flow cytometers. Currently, the statistical models for spectral unmixing remain the key limiting factor in ultra-high parameter panel design. In particular, increasing the number of fluorophores being unmixed reduces the tolerance to mismatch between the controls and the fully stained samples, as the visible spectrum of light is divided into smaller increments. Additionally, these skews in the data can be amplified by spreading inflation due to collinearity between dyes being used (Mage et al., 2025). Current commercial algorithms are limited in both error-correction features and their ability to account for heterogeneity of confounding autofluorescence within the sample. As such, “unmixing errors” are commonplace, to the extent that they are sometimes viewed as an inevitable part of least-squares unmixing approaches. These data artefacts are not, however, intrinsic to the data generated with current hardware, and are instead due to the application of insufficient statistical models during data processing and insufficient care during sample preparation. These limitations have the potential to be overcome, enhancing the data potential of spectral cytometry, through computational approaches that 1) error-correct the fluorophore signatures to refine the unmixing matrix, and 2) apply cell-specific autofluorescence extraction.

The criteria for producing ideal single-stained control samples are well established (Ferrer-Font et al., 2021; Roederer, 2002), but rarely followed in full. Some of problem lies with the lack of adequate software and reagent tools to allow users to correctly and cleanly extract spectral signatures. For example, we established with AutoSpill that simple intensity-based subtraction of fluorophore-negative from fluorophore-positive events in a single dimension produces less accurate spectra as it fails to exclude autofluorescent outlier events (Roca et al., 2021). Secondly, the selection of negative events is key to providing the correct intercept for the spectral signature vector (Roederer, 2002). The negative events should match the positive events in background (autofluorescence spectrum and magnitude), which is difficult or impossible to perform in available software for cell-based controls. This is no less a requirement when compensation beads are used, where the practice of selling “positive” beads admixed to chemically and spectrally distinct “negative” beads is widespread. A computational approach is therefore required to select “clean” events automatically from single-stained controls, in order to accurately measure the fluorophore spectra for unmixing.

A related computational challenge in current spectral unmixing approaches is the loss of resolution in the unmixed data due to cellular autofluorescence and fluorophore emission variability. This resolution loss can occur due to instrument noise, Poisson noise from fluorophore emission and variable autofluorescence noise from the cells or particles being measured (Roederer, 2016). All of these losses in resolution can be compounded by collinearity in the spectra of the dyes being unmixed (Kmet & Novo, 2025; Mage et al., 2025; Novo, 2022). A particular problem relates to the complex cellular mixtures that are prime candidates for analysis by spectral flow cytometry. Cellular autofluorescence results from a variety of factors, including metabolism, and thus varies by cell type, cellular state, and sample treatment. As such, errors in the unmixing due to misassignment of autofluorescence as fluorophore abundance insidiously introduce errors that can be as simple as inflation of the data (spreading) or population-related skews giving false positive or negative results.

To use spectral flow cytometry to its full potential, a data processing pipeline is needed that addresses these major shortcomings: the minimisation of unmixing errors through effective population selection and iterative refinement, the application of autofluorescence subtraction at the individual cell level, and the reduction of skewing and spreading through optimization of fluorophore signatures at the individual cell level. With AutoSpectral, we provide a tool to assist users in generating improved, reproducible unmixing from spectral flow cytometry data. AutoSpectral automatically selects appropriate events from single-stained controls, allowing the accurate calculation of the unmixing matrix, refined by iterative linear regression to minimise matrix error. We also provide a framework for identifying the major autofluorescence profiles within a cell mixture and applying the correct autofluorescence subtraction to each individual cell. Our computational pipeline uses the per-cell residual signal following application of the unmixing matrix to calculate the optimal autofluorescence subtraction, being the profile which minimises residual. The pipeline similarly identifies the variability in fluorophore spectra from the single-stained controls, and optimizes the unmixing matrix for each cell based on residual minimization. The aggregate impact of AutoSpectral is an error reduction frequently between 10- and 3000-fold on susceptible cell populations. AutoSpectral provides a single pipeline suitable for analysis of highly heterogenous cell populations.

## METHODS AND DATA

### Flow cytometry

Tissue data set: C57BL/6J mouse spleen, lung and brain tissues were prepared as previously described (Burton et al., 2024). The samples were stained with a set of 42 markers intended to cover the entire spectrum, deliberately placing markers in poor positions relative to known cellular autofluorescence contributions. Overnight staining was used (Whyte et al., 2022). The original data and additional details are available at Mendeley Data on DOI: 10.17632/y2zp5xx2hg.2. The AutoSpectral unmixed data are available at DOI: 10.17632/vzdxy8n7wf.1.

ID7000 data set: Human PBMCs were isolated from buffy coats. Cells were stained sequentially with a 40-colour optimized panel for surface markers and overnight for intracellular markers (Park et al., 2020; Whyte et al., 2022). Data were acquired on a 5-laser Sony ID7000. Data are available at Mendeley Data on DOI: 10.17632/2kvc98hc65.1.

OMIP-102: FACS Discover S8 data from OMIP-102 were kindly provided by Florian Mair. Details on the generation of these data are covered in the OMIP (Konecny et al., 2024). Versions of the data unmixed with AutoSpectral are available on Mendeley Data at DOI: 10.17632/j6yvfgdrcp.1.

Most figures were generated in R using ggplot2. Examples using unmixed data, with the exception of synthetic data sets, were plotted in FlowJo version 10.10.0 on Windows 11.

### Unmixing methods

AutoSpectral supports ordinary least squares (OLS), weighted least squares (WLS) and a version of Poisson-based unmixing using iteratively reweighted least squares. The latter is implemented in R via the MASS package using the glm call with the identity link function (in essence, each point’s observations become the weights for that point). An implementation is provided in C++ with an R wrapper via the AutoSpectralRcpp package.

For WLS unmixing, weighting in AutoSpectral is performed using detector noise measurements, such as the inverse of the detector coefficients of variation. For BD instruments, QSPE values encoded in the FCS file are used as measurements of detector reliability. For other cytometers, empirical estimates of the Poisson noise were obtained using mean fluorescence values across the entire sample, under the premise that the mean is the variance in a Poisson distribution.

### Data handling

Basic handling of FCS files in AutoSpectral occurs through *flowCore* 2.20.0, with *flowWorkspace* 4.20.0 for biexponential transformation to visualize the data. Graphics are largely via *ggplot2* 4.0.0, with *scattermore* 1.2 used for acceleration of pseudo-colour dot plots, and *grDevices* 4.5.1 and *viridis* 0.6.5 used for colour schemes (Wickham, 2016). Writing of newly unmixed FCS files is assisted by *AnnotatedDataFrame* in *Biobase* 2.68.0 (Huber et al., 2015). Reshaping of data is assisted by *dplyr* version 1.1.4 and *tidyr* 1.3.1 libraries. Parallel processing is implemented in R via futures using packages *future* 1.67.0 and *future.apply* 1.20.0. Rcpp is used to access C++, where processing is performed using RcppArmadillo and OpenMP.

AutoSpectral relies on automated gating developed in *AutoSpill* (Roca et al., 2021). Cell population boundaries are defined through density-based selection using kernel density estimation via *kde2d* in the *MASS* package version 7.3.65, as well as *interp.surface* from *fields* 17.1, *point.in.polygon* from *sp* 2.2.0*, convex.hull* and *tri.mesh* from *tripack* 1.3.9.3, and Voronoi tiling from *deldir* 2.0.4 (Venables et al., 2002). Details on tuning the automated gating are provided in articles associated with the AutoSpectral package. Unlike *AutoSpill*, pre-gated FCS files from third-party applications are also supported.

Large language models (LLM), particularly GPT4o, were used in the development of the code for AutoSpectral in situations where the output could be validated. Examples of LLM usage include identifying the correct axiomatic expression for ggplot, checking of loops and code for potential errors, conversion of *for* loops to vectorized versions, creation of initial C++ code from established R functions and the wrapping of functions into an R package using *roxygen*.

### Scatter-matching and removal of intrusive events

Scatter-matching adapts the automated gating approach. The *n* events with the highest signal in the peak detector for a given fluorophore are selected and a density-based boundary is defined using their forward and side scatter coordinates. This boundary is then applied to a matching unstained sample to select events with similar characteristics.

Autofluorescent event exclusion from single-colour control samples is performed using the corresponding unstained sample as a guide. The raw data from the unstained sample are scaled using the robust standard deviation and centred around the median in each detector channel. Principal components analysis is then used to identify the loadings driving the main sources of variation in the unstained sample, *i.e.*, the main autofluorescence spikes. These loadings are normalized and used as spectral signatures to unmix the unstained sample. Automated gating is performed on this low dimensional representation of the autofluorescence, identifying the main, dense population of cells near the origin as low autofluorescence. Cells outside this region constitute potentially intrusive autofluorescence. For each fluorophore, the corresponding unstained sample is assessed between the empirically identified peak signal in raw detector space for the intrusive autofluorescence and the prior-defined peak signal channel for the fluorophore. The trajectory of the main autofluorescence spike is identified using a robust linear model, and a boundary is established within two standard deviations of the predicted spike region. This exclusion boundary can then be applied to the matched stained control samples, allowing for exclusion of interfering autofluorescent events while retaining any signal from the fluorophore.

### Autofluorescence extraction

Single-cell autofluorescence extraction is handled initially through signature identification and pair-testing at the cellular level. Several autofluorescence signatures are identified from an unstained sample. The sample is unmixed into the fluorophore space to identify components of the autofluorescence most likely to affect the unmixed data. The combined unmixed and raw data are then mapped to a self-organizing map (SOM) (Kohonen, 1995; Van Gassen et al., 2015). Unlike FlowSOM, no metaclustering is performed, but rather the nodes themselves are used as the source of autofluorescence signatures. These autofluorescence signatures can then be tested pairwise, at a single cell level in combination with the optimized “best” fluorophore signatures to unmix the fully stained data, generating multiple OLS models of the data. The best model for each cell is selected based on the criterion of which model produces the lowest squared residual for that cell. In high parameter data sets where the residuals are already low, an alternative approach may be used, picking the model that produces the least absolute variance from zero in the fluorophore channels.

In the interest of reproducibility, all data relevant to the unmixing process, including the spectral signature mixing matrix, any weights used, the autofluorescence spectra and the per-cell assignments of which autofluorescence spectrum has been used are all encoded into the FCS files produced by AutoSpectral.

### Fluorophore spectral optimization

Single-cell fluorophore unmixing optimization is performed under the same framework as autofluorescence. Starting from single-stained control data, a SOM of the positive events is generated, and the background is subtracted from each node. This creates a set of potential variation in the spectral signatures of the fluorophore, as defined by the controls. To prevent contamination by autofluorescence signatures in rare populations, any spectra with cosine similarity less than 0.98 from the optimized single “best” spectrum for the fluorophore are excluded. These spectral variations are then used in the unmixing process as follows: First, the modelled autofluorescence component for each cell is subtracted from the raw data. This remaining data is then unmixed with all fluorophores in a single matrix (as is standard), using the single “best” spectrum for each fluorophore. Each cell thus starts with the “best” spectrum at baseline. From here, each cell is checked for whether it has detectable signal for each fluorophore, as defined by the 99.5^th^ percentile of the unstained sample. Each cell is then re-unmixed using only the fluorophores reaching the detection threshold, which reduces the unmixing space and increases the value of the residuals. For each fluorophore detected in a given cell, every variant spectrum for that fluorophore is tested pair-wise, keeping the spectrum that decreases the residuals. After all fluorophores have been processed and the optimal fluorophore variant signature assigned, the cell is unmixed again, this time using the selected spectral variants in the context of all fluorophores (not just those deemed present on the cell). Note that this differs from TRU-OLS, among other ways, in that this approach avoids discontinuities in the visualization and importantly allows the cytometrist to check for negative skews in the resulting unmixed data. The pure R implementation of this method is slow in use. In R, we therefore provide an approximation based on predicted changes in the raw component for a single endmember (fluorophore) without reconstructing the unmixing matrix, as this can be calculated cheaply in a matrix for all variants. We have implemented the exact method in C++ using OpenMP, accessible in R via Rcpp using the AutoSpectralRcpp package.

## RESULTS

### AutoSpectral improves spectral fitting in complex datasets

Current spectral unmixing approaches are limited by poor unmixing of fluorophore signal from cellular autofluorescence. To demonstrate these limitations, we used data from a 42-colour murine experiment produced on a 5-laser Cytek Aurora (**Figure 1**). These data were originally designed for a workshop on identifying and dealing with unmixing errors, and thus were intentionally difficult to unmix due to the marker:fluorophore combinations, in a realistic scenario. Unmixing artefacts apparent in this panel include unmixing-dependent spread, characterized as spreading and diagonal tilting of the unstained or negative data (e.g., the GFP channel versus BV510 in unstained, GFP-splenocytes), skewing of irrelevant fluorophore reporting in populations that are positive versus negative for a second fluorophore (e.g., correlated BV650 MFI in single-stained IgE-BV605+ cells), high spreading (e.g., high variance in single-stained CD19 BUV661+ cells versus APC), and diagonal spikes of autofluorescence (e.g., unstained samples from spleen, lung and brain) (**Figure 1A-D**).

**Figure 1.**
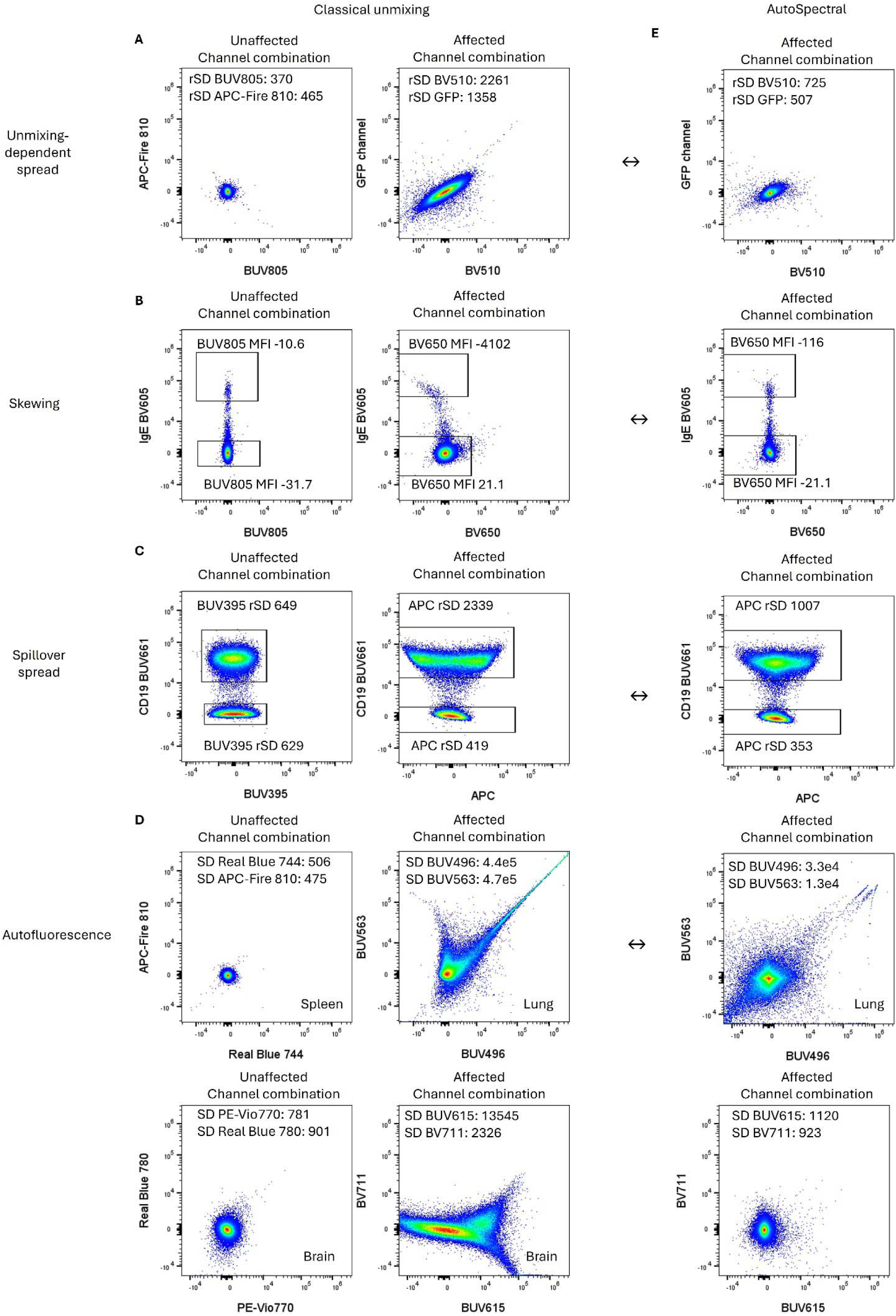
AutoSpectral addresses of common unmixing issues in high parameter spectral flow cytometry data. Spectral flow cytometry data from a 42-colour murine tissue experiment run on a 5-laser Cytek Aurora was analysed using classical approaches. Representative examples of fluorophore combinations that are unaffected or affected by **A)** unmixing-dependent spread, manifesting as tilting and bloating of the negative population (ungated unstained cells unmixed into the 42-colour space are shown), **B)** (anti-) correlation or skewing of populations in cells stained only with anti-IgE BV605, together with autofluorescence in the BV650 channel, and **C)** spillover spread, with trumpet-shaped distribution of BUV661+ events in a sample stained only with anti-CD19 BUV661. Gated on scatter signals. **D)** Multiple unadjusted autofluorescent signatures appear in the unstained tissue samples when unmixed with the fluorophore spectra. The magnitude and angle of the autofluorescent populations vary depending on the tissue source. Ungated data are shown. **E)** The same dataset was analysed using AutoSpectral, with double-headed arrows indicating the comparison plots under classical analysis. rSD: robust standard deviation. MFI: median fluorescence intensity.

To address these limitations of current spectral flow cytometry data processing pipelines, we adapted and expanded the AutoSpill compensation algorithm for spectral data, creating AutoSpectral. This approach is based on automated gating of events for the calculation of an initial spillover matrix and the use of robust linear regression to produce a fitted unmixing matrix (Roca et al., 2021). AutoSpectral was designed to improve the processing of spectral flow cytometry data through two independent stages. First, the extraction of fluorophore spectra from single-stained control cells is optimized through an autofluorescence-aware approach. These “clean” spectra can be used for any unmixing algorithm and may reduce skewing errors between similar fluorophores. Second, cellular heterogeneity is considered when unmixing the autofluorescence. In datasets with complex autofluorescence mixtures, this per-cell autofluorescence extraction reduces false positive and negative populations. Furthermore, since variability manifests as spread in unmixing, extracting autofluorescence on a per-cell basis has the effect of collapsing spread around zero between unmixed data for collinear fluorophores. Examples of the impact of the “AutoSpectral” unmixing approach on the unmixing problems are provided in **Figure 1E**.

The key initial point for extracting correct spectral profiles is to deal with the variability in cellular autofluorescence and remove the influence of noise (Roca et al., 2021; Roederer, 2002). The original AutoSpill package deals with this problem through automated removal of non-cellular events from single-stain controls and robust linear modelling (RLM) to de-weight noise (Roca et al., 2021). This gives an improvement over positive/negative population selection approaches, which fail to distinguish cells falling into the positive gate due to autofluorescence versus those reaching the positive threshold through fluorophore signal (**Figure 2A,B**), confounding the downstream calculations. By the simple means of assessing two-dimensions at a time, we can calculate the line of best fit along the fluorophore data. This process is facilitated by *a priori* knowledge of a detector where the signal to noise ratio is high (i.e., a “peak” detector), which can otherwise be distorted by the autofluorescence signal (**Figure 2C**). In robust modelling, points far from the line of best fit are iteratively de-weighted, reducing their influence on the calculated fit (**Figure 2D**). The coefficient of the fit, or in other words, the slope of the line, between the two detectors gives a spillover value for the dependent variable detector. This process is then repeated for each detector versus the “peak” detector, building up a “spillover” matrix, which is the spectral signature for the fluorophore. An advantage of the RLM approach is that the residual signal resembles cellular autofluorescence (cosine 0.88), demonstrating superior error-reduction (**Figure 2E-F**) and resulting in visible improvements to the unmixed data (**Figure 2G**).

**Figure 2.**
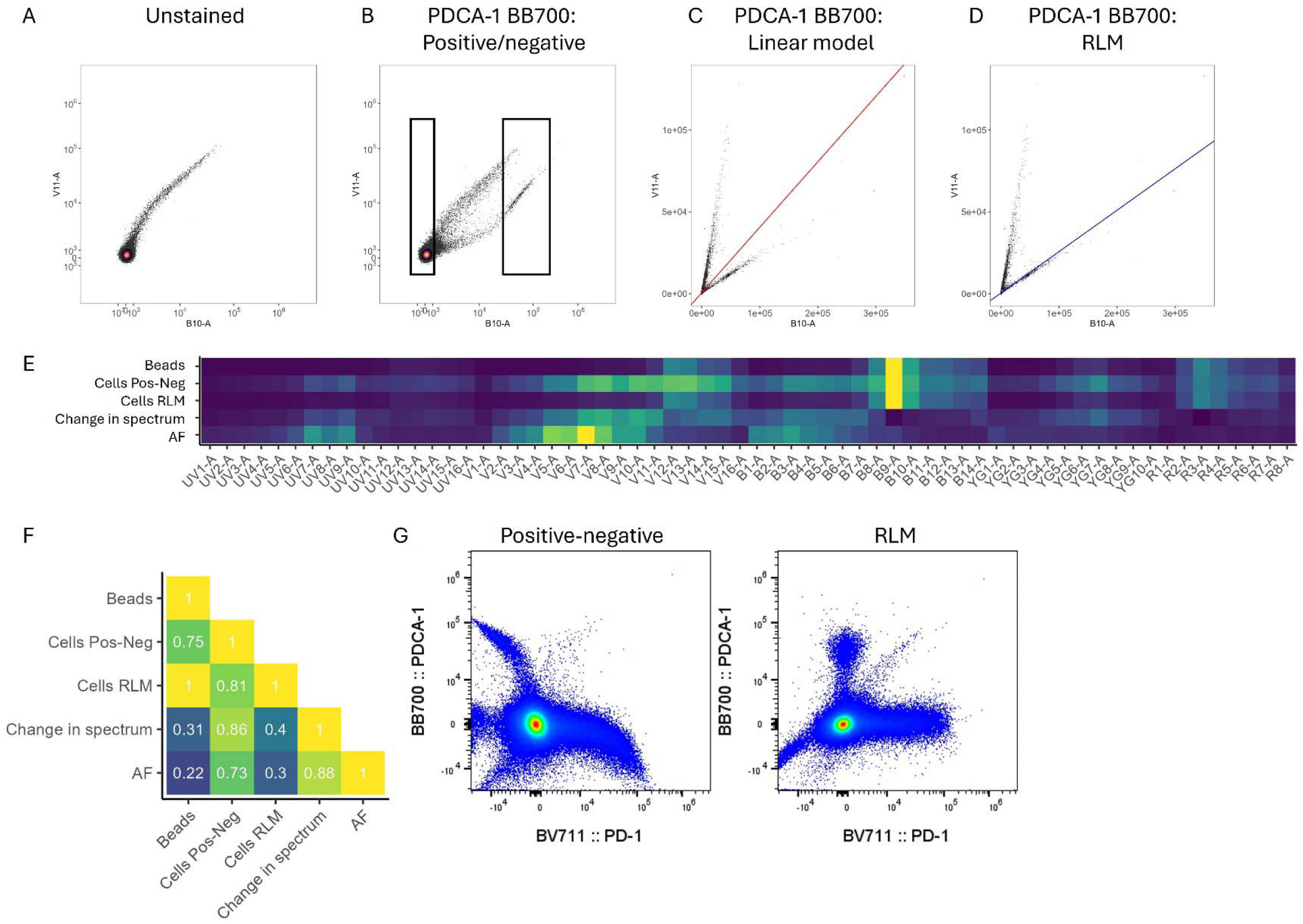
Robust linear modelling reduces the influence of autofluorescent outliers on spectral signature identification. The spillover fluorescence of murine splenocytes stained with anti-PDCA-1 BB700 was calculated based on classical positive-negative discrimination, linear models and robust linear models. **A)** Representative plot of unstained mouse splenocytes showing the autofluorescent events in the peak channel (B10-A) for BB700 versus a peak channel for autofluorescence (V7-A). Biexponential scale. **B)** Representative plot of mouse splenocytes stained with anti-PDCA-1 BB700 with a box showing a positive/negative gating approach, with both autofluorescence and BB700 fluorescence signals present. Biexponential scale. **C)** The anti-PDCA-1 BB700 control with the line of best fit from a linear regression (red) or **D)** a robust linear model (blue). Linear scale. **E)** Spectral heatmap showing the spectrum of BB700 as calculated from beads, from cells using the positive-negative gating approach, and from cells using the RLM approach. The impact of the cleaning steps on the identification of the BB700 signature using RLM approach is indicated through the change in spectrum achieved when shifting from the positive-negative gating approach to the RLM approach, by comparison with the autofluorescence present in unstained cells. The similarity of these profiles is calculated as **F)** the cosine similarity. **G)** Representative plots of OLS unmixed data with no autofluorescence extraction using either the positive-negative gating approach or the RLM approach to identifying the fluorophore signatures. Biexponential scale.

To further adapt the RLM approach to the context of spectral data, we added an additional step to match the background for fluorophore negative and positive events using scatter as a guide. This step improves unmixing accuracy, as the best results for unmixing are achieved using authentic single colour controls, however these controls can stain cell subsets with different baseline signal. For example, in a sample stained with a fluorescently conjugated anti-CD3 versus a sample stained with a fluorescently conjugated anti-CD14, different cell populations with distinct autofluorescent backgrounds are present in the brightest positive events. Accurate isolation of the spectral profile of the fluorophore without any involvement of differing cellular autofluorescence background requires the use of corresponding unstained cells matching the cells stained with the fluorophore (Roederer, 2002). While it may be impossible to match the events perfectly, we can match closely based on the scatter profiles of the positive (stained) cells. AutoSpectral does this without user input by first selecting the *n* (default is 500) events with highest intensity in the selected peak detector. This restriction of the calculations to the brightest events provides greater accuracy (Ferrer-Font et al., 2021; Roederer, 2002), and also speeds up the calculations. Once the positive events have been defined, a density-based boundary can be automatically generated on the forward and side scatter values for these events. Application of this boundary to a corresponding unstained sample allows for selection of events with matching forward and side scatter profiles, which corresponds to similar autofluorescence profiles (**Figure 3A,B**). We demonstrate that this process of scatter-matched negative selection reduces the influence of cellular autofluorescent background on the extracted fluorophore signatures (**Figure 3C,D**). Furthermore, the use of a separate unstained sample rather than internal negatives, which may carry low levels of fluorophore background, ensures the maximum separation between positive and negative events, improving accuracy (Roederer, 2002). Consequently, obvious unmixing errors are visibly reduced, often by orders of magnitude (**Figure 3E**).

**Figure 3.**
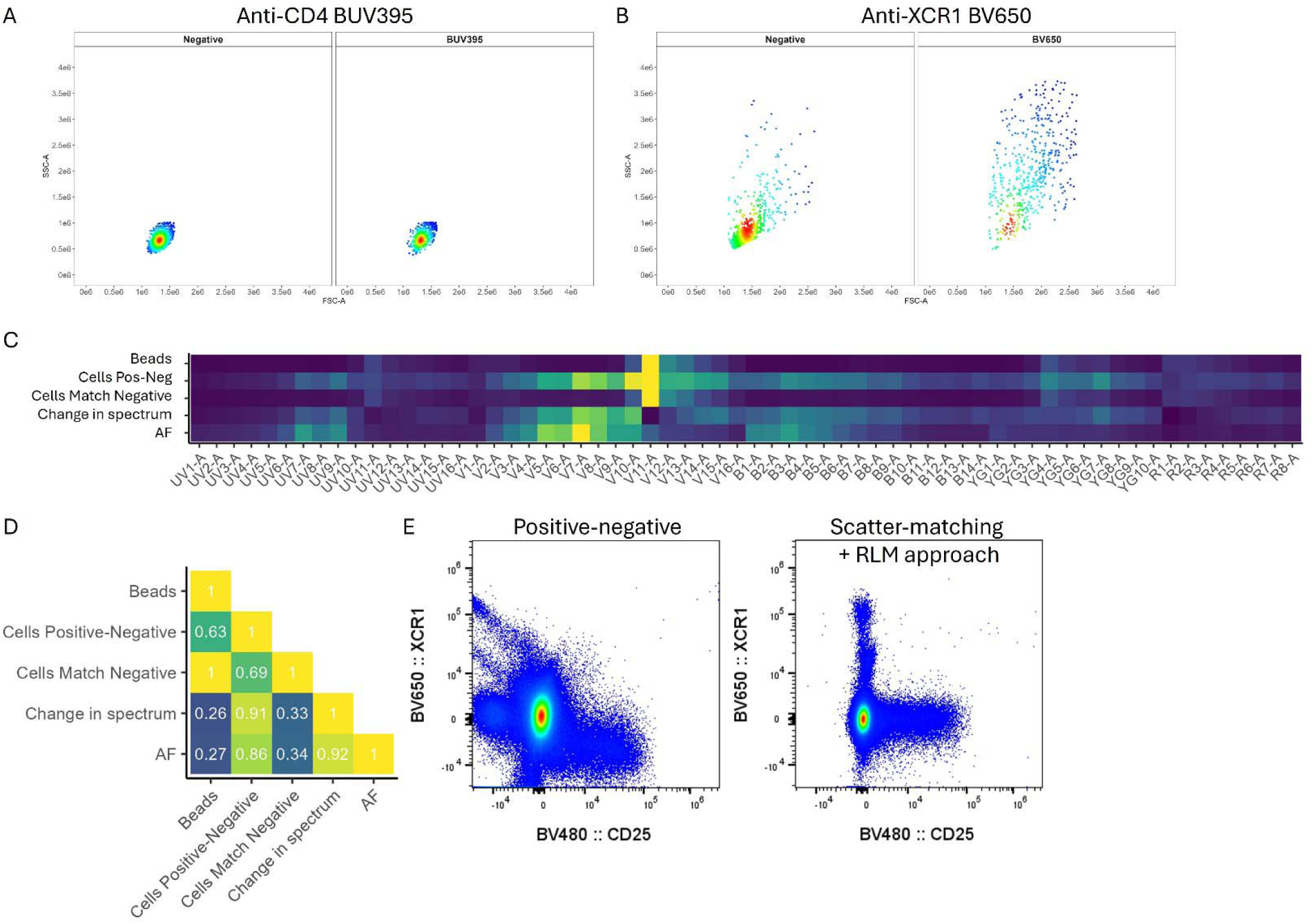
Scatter-matching negative and positive events aligns autofluorescence and reduces unmixing errors. **A)** Positive events following staining of mouse splenocytes with anti-CD4 BUV395, or **B)** anti-XCR1 BV650 were used to identify a negative population with matching forward and side scatter profiles. Representative scatter profiles of the gated positive population and the corresponding negative population selected by AutoSpectral. **C)** Spectral heatmap showing the spectrum of BV650 as calculated from beads, from cells using the positive-negative gating approach, and from cells using the scatter-matched approach. The impact of the cleaning steps on the identification of the BV650 signature using the scatter-matched approach is indicated through the change in spectrum achieved when shifting from unmatched to matched negative populations, by comparison with the autofluorescence present in unstained cells. The similarity of these profiles is calculated as **D)** the cosine similarity. **E)** Representative plots of OLS unmixed data with no autofluorescence extraction using either the unmatched or scatter-matched approaches to unmix.

Finally, we incorporated into AutoSpectral the removal of intrusive autofluorescence events when calculating the spillover matrix. AutoSpectral has the ability to deal with controls containing high ratios of noise to signal, however very high levels of autofluorescence in the form of intrusive events can distort results if the cells (e.g., macrophages, dead cells) are retained during the spectral signature calculation. By its nature, this autofluorescence is present in the unstained sample, and typically manifests as diagonal spikes in the data across multiple fluorescent detectors (**Figure 4A**). These spikes can be identified in an automated manner by scaling the unstained sample data according to the robust standard deviation per channel (to emphasize signals with high variance) and performing principal components analysis. The first two loadings from the principal components contain the highest amount of information regarding the directionality of the spikes of noise. These components are used as spectral signatures to unmix the unstained sample into a 2-dimensional representation of the autofluorescence noise space. On this representation, automated gating is applied to determine the standard-autofluorescence region around the origin (**Figure 4B**). When the peak signal detector for the noise population is plotted against the peak emission channel for the fluorophore, a boundary can be defined on the unstained sample, identifying the intrusive events (**Figure 4C**). This boundary can then be applied to the single-stained sample, excluding events with the boundary from consideration in calculation of the fluorophore spectrum (**Figure 4D**). Exclusion of noisy events is only applied to stained controls, thus permitting scatter-matching in the event that the positive events in the stained control are biologically identical to the noise in the unstained. This autofluorescence identification and exclusion process reduces collinearity between the spectra calculated using messy single-stained cell controls (**Figure 4E,F**). Notably, the intrusive event exclusion is only performed on the single colour controls to determine the spectral unmixing, and thus the visible improvement in unmixed samples is due to reduced unmixing errors rather than the removal of any events from the experimental data (**Figure 4G**).

**Figure 4.**
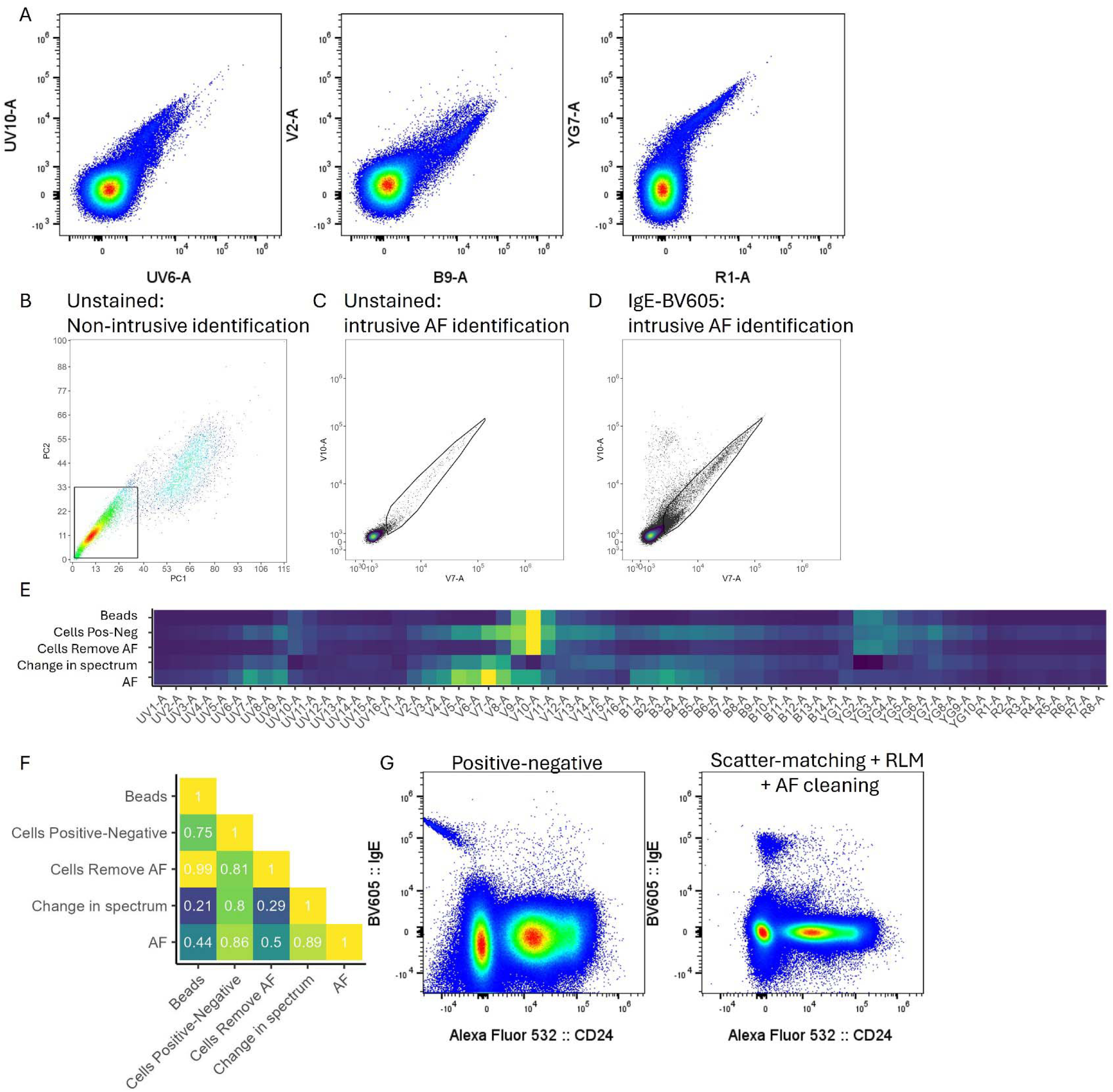
Removal of intrusive autofluorescent events improves spectral determination. **A)** Representative plots showing examples of intrusive autofluorescence spikes across multiple detectors in spectral flow cytometry data. Unstained mouse splenocytes. **B)** Representative example of gating non-intrusive cells on an unstained cell sample unmixed using components identified by PCA. **C)** A spline-fitted gate around the potentially intrusive autofluorescent events, using the lower bound established in A. The gate is created on the peak channel for the cells outside the region in A versus the a priori-known peak channel for the fluorophore to be used, in this case, BV605. **D)** Representation of the intrusive event gate on the single-stained anti-IgE BV605 control, indicating cells to be excluded during the unmixing calculation. **E)** Spectral heatmap showing the spectrum of BV605 as calculated from beads, from cells using the positive-negative gating approach, and from cells using the removal of intrusive events. The impact of the cleaning steps on the identification of the BV605 signature using intrusive event removal is indicated through the change in spectrum achieved if intrusive events are removed, by comparison with the autofluorescence present in unstained cells. The similarity of these profiles is calculated as **F)** the cosine similarity. **G)** Representative plots of OLS unmixed data with no autofluorescence extraction with and without intrusive event subtraction prior to calculation of the unmixing matrix.

### AutoSpectral enables simultaneous spectral fitting of autofluorescence in mixed cellular populations

Spectral flow cytometry allows handling of cellular autofluorescence through extraction of the fluorescent signature(s) present in the unstained cell samples alongside fluorophore signals. When paired with an accurate autofluorescence signature, this can reduce background and false positivity in the unmixed data. A limitation of current algorithmic approaches is that they do not accurately account for the inherent variability of cellular autofluorescence in both magnitude and profile, leading to spectrum analyses that are, at best, suitable for only one profile within the heterogenous mixture. Iterative manual extraction of multiple autofluorescence parameters has been proposed as a solution (Jameson et al., 2022; Niewold et al., 2020; Pilkington, 2024; Roet et al., 2024; Wanner et al., 2022), however these processes are time-consuming and require assessment of an unknown quantity of options in permutations which are often ill-conditioned without well-defined endpoints or metrics. To implement this approach in a systematic and automated manner, we designed AutoSpectral to include an identification step whereby the underlying autofluorescence signature of each individual cell is determined, allowing accurate autofluorescence extraction and improved implementation of the spectral unmixing of signals derived from fluorophores. The approach is based on using the unstained data to identify the diverse autofluorescence signatures present in the mixture, and then testing the appropriateness of each signature on each cell within the sample, using the residuals from the least-squares unmixing calculation. As the residuals are the divergence between the unmixing model fitted data and the original observations, the signature that results in the lowest residuals is the one where the fluorophore signature has the best fit, resulting in superior data fitting.

To test the validity of this approach, we generated synthetic data based on the mouse lung, which is well known to contain multiple highly divergent autofluorescent populations at relatively high frequency (**Supplementary Figure 1**). When mapped to a SOM, the full diversity of spectra present in the unstained lung sample were visualized as normalized fluorescence traces (**Supplementary Figure 1A**). Using the pairwise autofluorescence testing, the “best fit” autofluorescence was assigned to each cell as a parameter called “AF Index”. Plotting the unstained mouse lung data in detectors that receive high levels of autofluorescent signals reveals neatly discriminated populations, corresponding to these AF indices (**Supplementary Figure 1B**). The application of these AF indices leaves only modest spread even in 42-colour-stained samples (**Supplementary Figure 1C**). The effectiveness of this approach is similar to that achieved through an alternative, in which the selection of which autofluorescence spectrum to use per-cell is based on reducing the absolute values of fluorophores signals rather than minimising residuals (**Supplementary Figure 1D**). This method operates independently of the residuals, and thus can outperform residuals-based selection when panels become larger. Both methods match autofluorescence signatures to cells effectively, as evidenced by high cosine similarity values between the selected autofluorescence spectrum and the cell’s underlying signature (**Supplementary Figure 1E-G**). To test the validity of this method, we generated synthetic data with known autofluorescence and fluorophore contributions, using real lung unstained data and adding simulated fluorophore signals in the 42-dimensions of the corresponding panel. When tested against single autofluorescence extraction using a median spectrum, the per-cell autofluorescence unmixing model reduced the error in identifying both the autofluorescence and fluorophore components of the synthetic data (**Supplementary Figure 1H-I**). Autofluorescence spectra selections were essentially unaffected by the presence of the synthetic fluorophore layer, demonstrating the power and reproducibility of this method (**Supplementary Figure 1J**). Furthermore, the identified associations of autofluorescence spectra and cells are likely biologically meaningful as the generated “AF Index” parameter shows associations with scatter signals (**Supplementary Figure 1K**) and, in the fully stained data, with distinct cell types (**Supplementary Figure 1L**).

We next tested this approach on real-world complex stain sets. Using a complex stain set performed on mouse lung, brain and spleen samples (**Figure 5A-F**), per-cell autofluorescence extraction correctly reassigned the fluorescence signal in the unstained samples to the autofluorescence channel (**Figure 5A,C,E**). For the corresponding fully stained samples, per-cell autofluorescence extraction removed features associated with artefacts, including strongly correlated signals (**Figure 5B,D,F**). In particular, these changes corresponded to corrections in known biological errors, including the shift among autofluorescent F4/80^low^ microglia to misallocate signal to a T cell-driven genetic reporter (**Figure 5B**), the misallocation of signal within lung CD11c^hi^ cells (alveolar macrophages/eosinophils) to IgD, which is restricted to the B cell lineage (**Figure 5D**), and the false signal of the CD4 T cell marker on splenic F4/80^+^ macrophages (**Figure 5F**). When compared to existing autofluorescence extraction methods, either single or multiple, per-cell extraction correctly re-assigned unmixed values for cells appearing as diagonal spikes, negative skews, positive skews and off-origin negatives, without creating unmixing-dependent spread (**Figure 5G**). Visually, we observed improved resolution in a 40-colour human PBMC panel run on the SONY ID7000 (**Figure 5H**) and on the 50-colour OMIP-102 data from the BD FACSDiscoverS8 (**Figure 5I**). Quantification of the resolution using robust standard deviation on unstained samples showed a decrease in variance in all cases, particularly in channels with the highest variance, which are those most affected by autofluorescence and unmixing-dependent spread (**Figure 5J**). We understand this to be consistent with autofluorescence being a form of improperly corrected noise in the OLS unmixing model, and it being amplified by collinearity of fluorophore spectra (Kmet & Novo, 2025; Mage et al., 2025). Finally, we demonstrate that per-cell autofluorescence extraction improves the accuracy of the unmixing model, as assessed by the reduction in absolute value of the residuals (**Figure 5K**).

**Figure 5.**
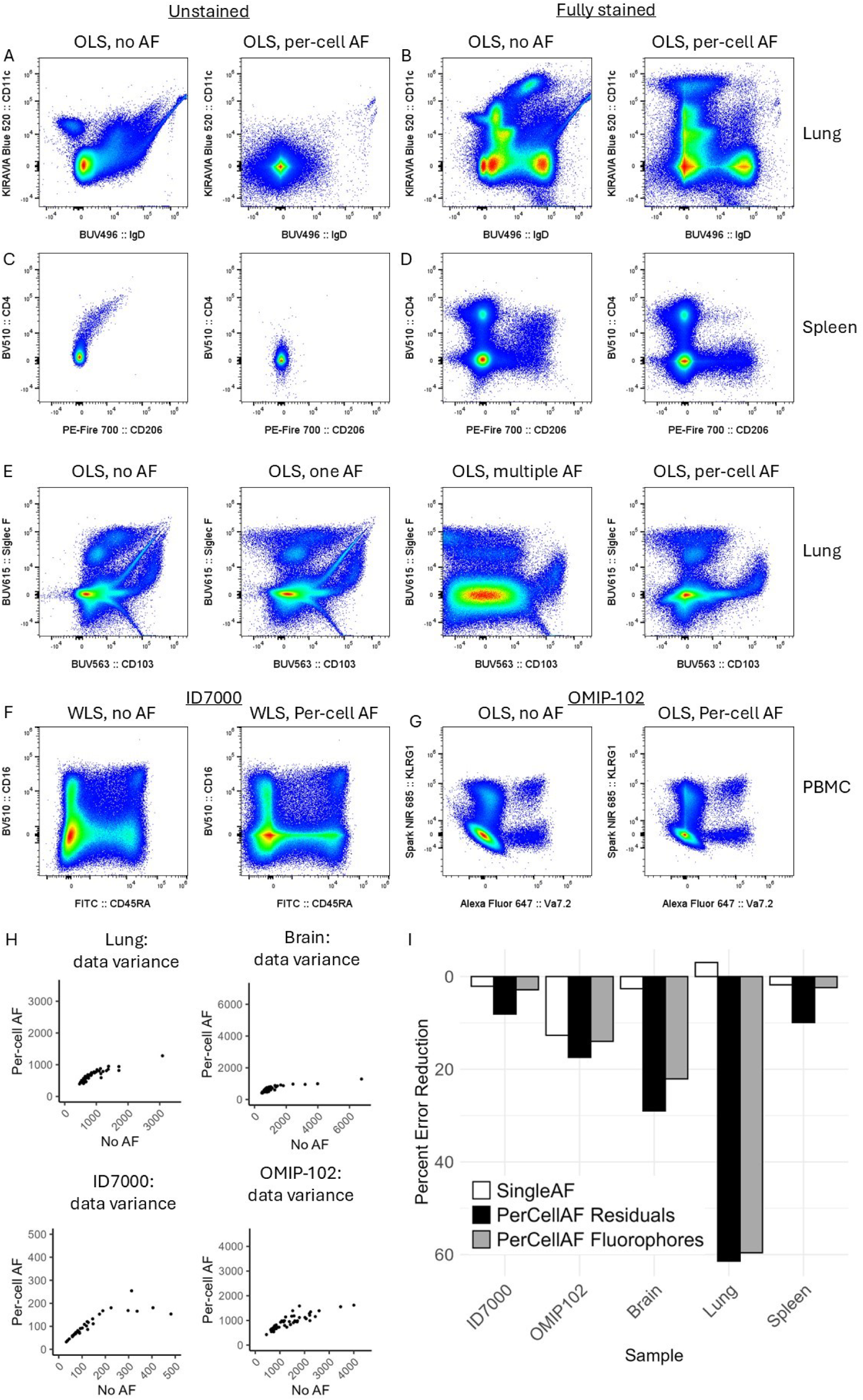
Per-cell autofluorescence extraction improves unmixing model accuracy and reduces spread. Using a 42-colour mouse panel, the spillover matrix was calculated using either the OLS approach alone, or the OLS approach combined with AF matching performed at the per-cell level. Comparison between the two approaches is shown for **A)** ungated events from the unstained sample of mouse lung, **B)** viable CD45+ gated events from the stained sample of mouse lung, **C)** ungated events from the unstained sample of mouse spleen, or **D)** viable CD45+ gated events from the stained sample of mouse spleen. **E)** The effect of AF matching on unmixing of 42-colour-stained mouse lung, with the analysis performed without AF extraction, with a single median AF spectrum, with multiple AF spectra or with per-cell AF extraction. Gated on viable CD45+ cells. **F)** Representative plots from the 40-colour ID7000 human PBMC data set, with and without per-cell AF extraction, and from the **G)** 50-colour OMIP-102 human PBMC data set, with and without per-cell AF extraction. **H)** Impact of per-cell AF extraction on unmixed data variance (per-channel mean absolute deviation) on unstained samples. Unstained samples from each data set were unmixed using OLS (or WLS for ID7000 data) either without any AF extraction “No AF” or per-cell AF extraction “Per-cell AF”. Scatterplot of data variance for individual unmixed fluorophore channels, under either the “no AF” or “Per-cell AF” method. **I)** Percent reduction in error achieved by different approaches of AF extraction, as calculated by the difference in absolute residuals compared to unmixing without AF extraction.

### AutoSpectral enables simultaneous spectral fitting of fluorophore signals on a per-cell basis

Variability exists in fluorophore emissions and measurements thereof on a per-cell basis, with deviation around the median spectrum manifesting as spillover spread. This spread can propagate through collinear channels, creating a loss of resolution. To study this, we used BUV661, an ultraviolet-excitable tandem dye with an acceptor moiety directing emission into the red range of visible light. Among cells stained with anti-CD19 BUV661, there is noticeable variation in the relative emission detected in red laser channels (**Figure 6A**). When unmixed using OLS in the context of the 42-colour panel containing APC and other red laser-excited dyes, the BUV661-stained cells exhibit variability with respect to red laser dyes, with the characteristic trumpet shape when viewed on a biexponential transformation (**Figure 1C** and data not shown) (Roederer, 2016). This variability can lead to false “double positive” events, such as CD19 expression spilling into the Foxp3 gate, despite no biological co-expression (**Figure 6B**). In this case, the spillover spread is not driven by autofluorescence, and therefore not markedly reduced by per-cell autofluorescence extraction, although the minor extraneous signals are (data not shown). When the BUV661 spectrum is allowed to vary on a per-cell basis between the range of valid options available from the single-colour control and a best fit per-cell is selected based on residual minimization, the variability in APC values for BUV661^+^ events diminishes substantially, reducing the number of false “double positive” events (**Figure 6B**). This improvement also holds up in synthetic datasets where ground truth is known, and the fluorophore optimization can be shown to improve the matching of each cell’s underlying fluorophore signature (**Supplementary Figure 2**). An additional, non-trivial benefit of per-cell fluorophore fitting is the reduction in skews or non-biologically correlations in the data, often termed “unmixing errors”. These errors can result from mismatches between the defined spectra from the controls and the actual spectra present in the fully stained samples.

**Figure 6.**
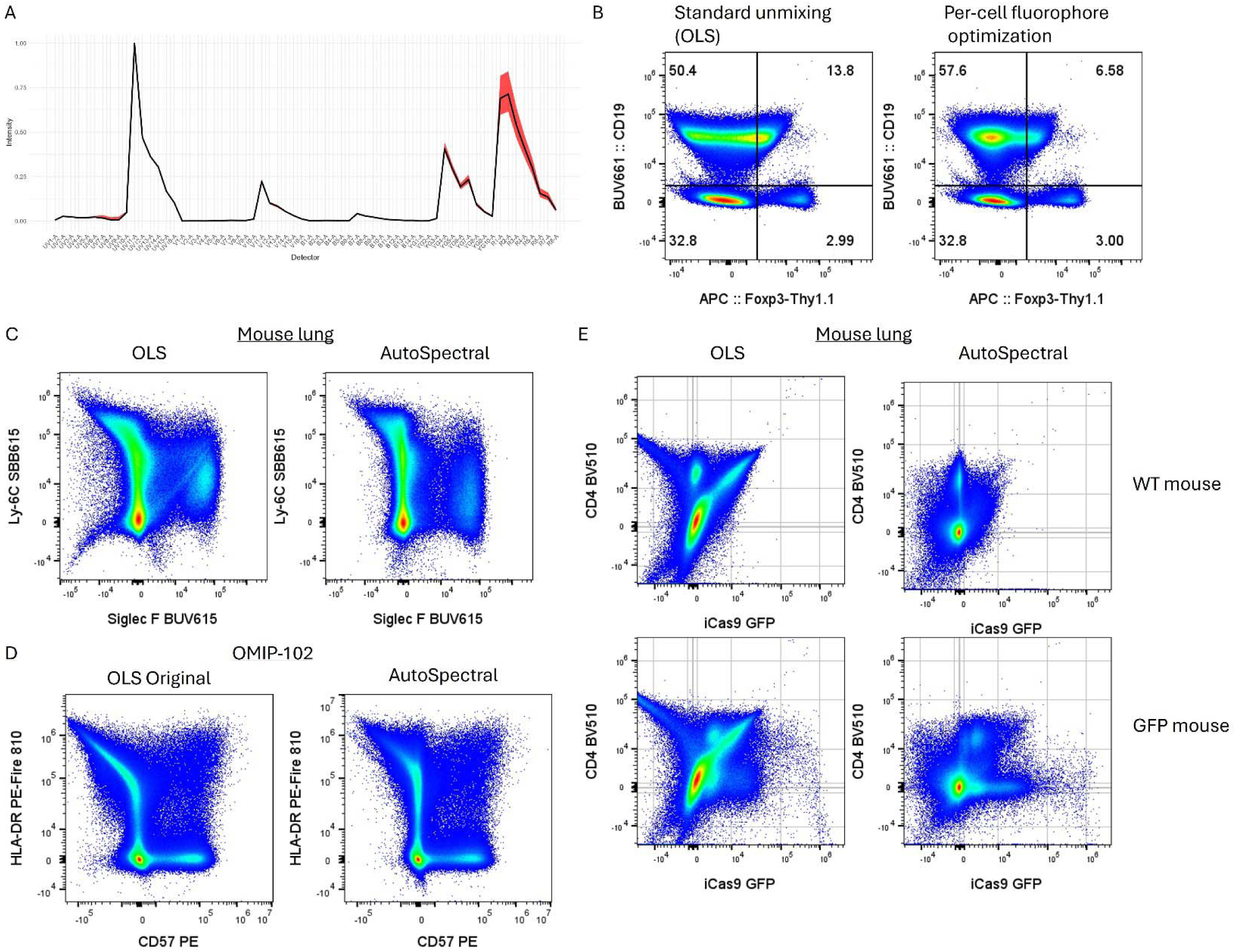
Per-cell fluorophore optimization reduces spillover spread and reduces data skewing. **A)** Spectrum for BUV661 obtained from murine splenocyte stained with anti-CD19 BUV661, with median signal indicated by solid black line, and signal range indicated in red. **B)** The effect of per-cell fluorophore optimisation is shown using representative plot of mouse splenocytes from the 42-color panel, gated on viable single CD45^+^ cells. Double positive events are not expected biologically and represent either spread or non-specific staining. **C)** Unmixing skew for SBB615 versus BUV615 with OLS or per-cell (AutoSpectral) optimization in the mouse 42-colour dataset. **D)** Unmixing skew for PE/Fire 810 versus PE in the OMIP-102 data, OLS or per-cell optimization. **E)** Detection of weak GFP signals in the mouse lung, OLS vs per-cell optimization. Data are shown from a wild-type (“WT”) mouse that does not express GFP and a CD4Cre iCas9-GFP (“GFP”) mouse that expresses GFP within the CD4+ and CD8+ populations. Data in B-E are shown gating on single leukocytes (lymphocytes, monocyte and granulocytes) by scatter without any further clean-up.

The spectral fitting of fluorophore data via AutoSpectral enables the distribution of spectra from the single-stained controls to be fitted to the spectrum of each cell in the stained data, allowing for improved data fitting. To test the ability of per-cell spectral optimization to partially correct negative skews in the data, we used several real-world examples of this phenomenon. First, for the 42-colour panel, high expression for Ly-6C in the mouse lung drives a SBB615 signal that is associated with negative skewing in the BUV615 channel (**Figure 6C**). Second, for OMIP-102, PE/Fire 810, a dye prone to tandem breakdown, appears to show more breakdown and lower expression in the single-colour control than in the fully stained samples, manifesting as a non-biological negative correlation between HLA-DR PE/Fire 810 and CD57 PE in the samples (**Figure 6D**). In both cases, when the best fit for each fluorophore, based on the observed range present in the single colour control, is selected at the level of the individual cell, these skews are noticeably reduced (**Figure 6C,D**).

Finally, we tested the collective impact of the combined AutoSpectral pipeline, using a challenging real-world example where the ground truth state is known. Here we turned to the mouse tissue data set, using the example of a weak transgenic reporter for GFP present in T cells, which results in poor resolution in the lung. Using a wildtype mouse as ground truth for negative GFP expression, classical approaches can detect the subtle shifts in a CD4+ and CD8+ population, however these shifts are obscured by artefacts (**Figure 6E**). The use of AutoSpectral on this dataset reduces spread, removes diagonal spikes and negative skews, substantially cleaning up the wildtype plot and allowing the weak GFP+ populations to be clearly discriminated in the GFP transgenic mouse (**Figure 6E**). Together, these improvements demonstrated the ability of AutoSpectral to substantially improve the data quality of complex spectral flow cytometry experiments, even with weak fluorophores in problematic combinations and challenging tissues.

## DISCUSSION

AutoSpectral provides a full automated workflow for optimizing unmixing of spectral flow cytometry data. The innovations we present address intrinsic variability present in spectral flow cytometry data. This variation is present even within the single-stained controls, needed to define the baseline spectra for each fluorophore. While ideally uniform, these controls are inherently multi-coloured and both qualitatively and quantitatively variable. Even under simplistic settings such as supposedly uniform cell line cultures, differences in viability or metabolism can alter the fluorescent background, creating a variable mixture. Ideal cellular controls are typically based on complex mixtures of cell types, with markedly different fluorescent backgrounds. In controls with high intensity “bright” staining that is evenly distributed across the heterogenous population, such noise tends to be inconsequential in regard to correctly defining a median “optimal” signature. More commonly, markers are expressed at low or variable levels, with brightness levels that overlap with the tail of the cellular background noise distribution, and expression is biased towards particular cell subsets, creating a background autofluorescence signal that differs between the “positive” and “negative” events. This leads to imprecision in distinguishing the fluorophore background from the autofluorescence background, creating the suboptimal unmixing matrices which led to artefacts in the unmixed data. Compensation beads with supposedly uniform background profiles were introduced to circumvent this issue and are quite popular due to their relative ease of use. Bead-based controls often fail, however, to recapitulate the exact spectral profile of the fluorophores as seen on the stained cells (Ferrer-Font et al., 2021; Konecny et al., 2024; Liechti et al., 2023; Park et al., 2020; Shevchenko et al., 2024), and fail to adequately account for cellular autofluorescence. Thus, while single colour beads are a functional workaround, the ideal solution for spillover calculation requires a statistical model that can accurately dissect signal origins from single colour-stained cells.

The essence of cytometry is variability at the cellular level. The mathematics of classical unmixing approaches, however, relies on fitting the data to what are effectively median measurements of a mixed population. At best, this creates distributions of error in the data around the median (spreading), and in cases where the fully stained samples differ from the controls, the distributions may be skewed, creating visual unmixing errors.

Definition of the cellular and fluorophore spectra can be derived from the use unstained and single colour cells, creating a signature matrix permitting individual autofluorescence and fluorophore signals to be extracted from the actual samples. For highly heterogeneous and complex samples, classical unmixing is well appreciated to fail to produce unmixing matrices that are invalid for multiple cell types within the mixture. Skilled analysts can manually clean control populations (Ferrer-Font et al., 2021) and create multiple unmixing matrices for different cell or sample types, such as is now widely used to account for variable autofluorescence spectra (Kharraz et al., 2022; Niewold et al., 2020; Pilkington, 2024; Roet et al., 2024; Wanner et al., 2022).

These approaches, however, have intrinsic limitations compared to algorithmic-based solutions.

Due to the previous absence of a systematic algorithm capable of effectively extracting multiple autofluorescent signatures from a mixed sample, most manual solutions to unmixing in complex samples involve a “guess and check” approach. This approach requires the researcher to sort through the high dimensional data from the unstained sample(s) for “populations”, extracting signatures from these and adding them in various permutations to the unmixing matrix (Ferrer-Font et al., 2021; Wanner et al., 2022).

While this practice can be improved through the use of clustering to identify potential populations (Jameson et al., 2022; Roet et al., 2024), this does not solve the issue of identifying which signatures(s) to use (Pilkington, 2024). Overall, while previously necessary, these approaches are laborious, require a high level of expertise, are prone to becoming poorly conditioned and reducing resolution (Ferrer-Font et al., 2021; Jameson et al., 2022; Pilkington, 2024; Roet et al., 2024). The very nature of this process makes it nearly impossible to replicate, even when starting from the same set of control samples. AutoSpectral provides a mathematical solution to the problem, enabling the process of both control cleaning and application of the appropriate autofluorescence signature to be automated. The statistical modelling approach also enables variation in fluorophore spectra to be similarly accounted for, an issue which, while mathematically identical, is too nuanced to permit manual approaches. We also wish to highlight the reproducibility of AutoSpectral: the mathematical basis of the approach allows consistent results to be obtained, with lower error rates, from the same set of data. As the approach is deterministic, the same outcome is achieved regardless of the user or run, making it an appropriate choice for analyses where reproducibility is prioritised.

We believe this to be the first published approach to manage both cellular autofluorescence and fluorescence spectra on a per-cell level. We note, however, that the proprietary algorithm Ozette Resolve appears to deal with unmixing on a per-cell basis and may perform spectral optimization accordingly. The unmixing approach in that model is not publicly available, precluding direct comparison, however it appears to differ in its underlying mathematical model. Among published works, the closest approach we are aware of is the recent article by Kmet and Novo, which drops irrelevant fluorophore spectra from the mixing matrix cell-by-cell to reduce spread (Kmet & Novo, 2025). This TRU-OLS approach is distinct from AutoSpectral in that it deals with the propagation of the noise driving spread rather than the underlying cause, but it is compatible with AutoSpectral and could be integrated.

In this work, we demonstrate that the best fit autofluorescence assignments for cells correlate with scatter profiles, biological cell types and imaging parameters. We suggest, as previously noted (Pilkington, 2024), that autofluorescence provides meaningful information regarding cell states and point out that label-free identification of cell types based on intrinsic fluorescence is used by multiple platforms (Kinegawa et al., 2023; Ota et al., 2018; Suzuki et al., 2019). To make the most of this information, the best fit cell-specific autofluorescence assignments are encoded into the FCS files produced by AutoSpectral as an additional parameter called “AF Index”, which takes the form of numbers 0-n, where 0 indicates that no autofluorescence extraction was the best fit. We expect that users may find this information relevant when assessing cellular phenotypes and may allow for biological discovery in populations with limited availability of fluorescent biomarker reagents. While we have restricted AutoSpectral to the use of fluorescence detector channels for the identification of cell-specific autofluorescence profiles, it is likely that inclusion of imaging parameters, where available on image cytometers, would enhance the identification process and provide new biological insights.

### Limitations of the approach

While AutoSpectral can reduce unmixing error, improving the assigned cell positions of error-prone populations by up to 9000-fold, the ability to calculate the unmixing matrix is dependent on high quality data. In particular, AutoSpectral does not reduce the need for high quality controls, and cannot be used as a substitute for suboptimal selection of experimental controls or poor sample preparation or data collection. Indeed, the optimal use of AutoSpectral requires enhanced diligence in the preparation of control and samples, to provide the quality input data needed for the unmixing calculations.

The repeated unmixing process used by AutoSpectral to optimize the modelling is also slower than a single-shot OLS unmixing. The impact of this is limited, however due to the high optimisation of linear algebra algorithms, with the rate-limiting factor in unmixing process often being the overall handling of FCS files, particularly in R. For the autofluorescence portion, AutoSpectral calculates all events simultaneously and merely picks the best per-cell result, rather than performing a single unmixing per cell. If timing is considered against the current best approach of manual multiple autofluorescence extraction, for which there are astronomical permutations, AutoSpectral is exceedingly quick. We have implemented the per-cell fluorophore spectral optimization in C++ with parallelization, and in addition to the exact calculation method, we provide an approximation using Woodbury-Sherman-Morrison rank-one updating (Hager, 1989; Sherman, 1949). With the latter approach, the unmixing calculation for the fully stained lung sample used here can be performed on a mid-range Dell laptop with a 3.00GHz i7 core in approximately two minutes, or in other words, a similar amount of time as is required for the acquisition on the cytometer. As such, the total calculation time is compatible with downstream data analysis. Currently, however, most while aspects of the AutoSpectral pipeline would improve the calculation of unmixing matrices for use on sorting machines, the full implementation may not be feasible for the real-time calculations required when sorting. For the application of per-cell autofluorescence or fluorophore calculations, an approximate implementation in real-time sorting may, however, be achievable through machine learning models, providing much of the benefit observed here.

## CONFLICT OF INTEREST STATEMENT

Oliver Burton provides flow cytometry consulting services and has consulted for Bio-Rad, makers of StarBright dyes. The other authors declare no conflict of interest.

## DATA AVAILABILITY STATEMENT

Data files have been deposited on Mendeley Data. Additional guidance for effective use of AutoSpectral is provided on GitHub in markdown format and on https://www.colibri-cytometry.com/blog. The code for AutoSpectral is available at https://github.com/DrCytometer/AutoSpectral. A faster version for per-cell unmixing is available at https://github.com/DrCytometer/AutoSpectralRcpp.

## ACKNOWLEDGEMENTS

The authors gratefully acknowledge the Department of Pathology flow cytometry facility from the School of the Biological Sciences at the University of Cambridge as well as the FACS core at the University of Leuven for their support and assistance in this work. We wish to thank Florian Mair for providing data from OMIP-102. We thank Charlotte Christie-Petersen for providing a set of data for the NovoCyte Opteon. We thank Jochen Lamote for insightful discussion regarding imaging data and autofluorescence. This work was supported by the Medical Research Council grant MR/Y004450/1 through the MRC Proactive Vaccinology consortium IMMPROVE (https://immprove.ac.uk/), and the Wellcome Trust through Wellcome Investigator Award 222442/A/21/Z.

## Notes

https://github.com/DrCytometer/AutoSpectral

