## Supplementary material for "AutoSpectral improves spectral flow cytometry accuracy through optimised spectral unmixing and autofluorescence-matching at the cellular level"

**SUPPLEMENTARY FIGURE LEGENDS:**

**Supplementary Figure 1. Principles of single cell autofluorescence identification and extraction**. **A)** Autofluorescence spectra obtained from mouse lung in AutoSpectral using SOM clustering. **B)** Representative flow cytometry plot of unstained mouse lung raw data without autofluorescence extraction, with points coloured by AutoSpectral autofluorescence assignment (“AF Index”). The plotted channels were selected at peaks of the autofluorescence spectra in A. **C)** Unmixing following testing of each cell against each autofluorescence spectrum observed in the unstained sample (A), with selection based on best minimization of the absolute residual per cell or **D)** based on the best minimization of the fluorophore signal per cell. **E)** Distribution of cosine similarity values between the various lung autofluorescence spectra, mimicking random assignment of AF Index. **F)** Distribution of cosine similarity values between the unstained cell’s spectrum and the assigned AF spectrum, with assignment by residuals or **G**) by minimization of fluorophore signal. Statistics by Komolgorov-Smirnov (KS) test versus F. **H)** Distribution of errors in the prediction of autofluorescence data for 10000 mouse lung cells using a single median AF signature or per-cell AF unmixing. Predictions are shown with and without synthetic fluorophore data added to unstained mouse lung data. Data were generated using unstained mouse lung raw data. These raw data represent autofluorescence ground truth. The data were then unmixed using OLS with a single median AF signature or per-cell AF extraction. These unmixing models were then used to revert the unmixed data back to the original raw detector space in a prediction of the original data. Each cell’s error was calculated as the sum of absolute differences between the model’s prediction and the original data. Each cell’s error is plotted as point on the graph, with a violin in overlay to assist visualization of the distribution. Additionally, synthetic data was generated by adding a layer of in silico-generated random fluorophore signals (derived from the 42-color fluorophore spectral matrix) onto the unstained sample’s raw data, creating a situation mimicking fully stained data, but with the underlying unstained cell’s observations being known as ground truth. Errors for these synthetic samples were calculated using the same process. **I)** Distribution of errors in the unmixing of the synthetic fluorophore data when real mouse lung cell autofluorescence is added, either with a single median autofluorescence signature or with per-cell AF selection. Statistical testing by KS test. **J)** Distribution of cosine similarities in AF spectrum assignment with and without synthetic fluorophore data added to unstained mouse lung data. A value of 1 indicates identical spectral assignments. **K)** Side scatter distributions by AF Index. Cells were grouped by the AF Index (autofluorescence signature) assigned by AutoSpectral and the side scatter values (unused in the AF assignment) were plotted. **L)** Distribution of AF Indices by cell type for mouse lung CD8 T cells, CD19 B cells, CD11b Ly-6C monocytes and CD11c Siglec F alveolar macrophages.

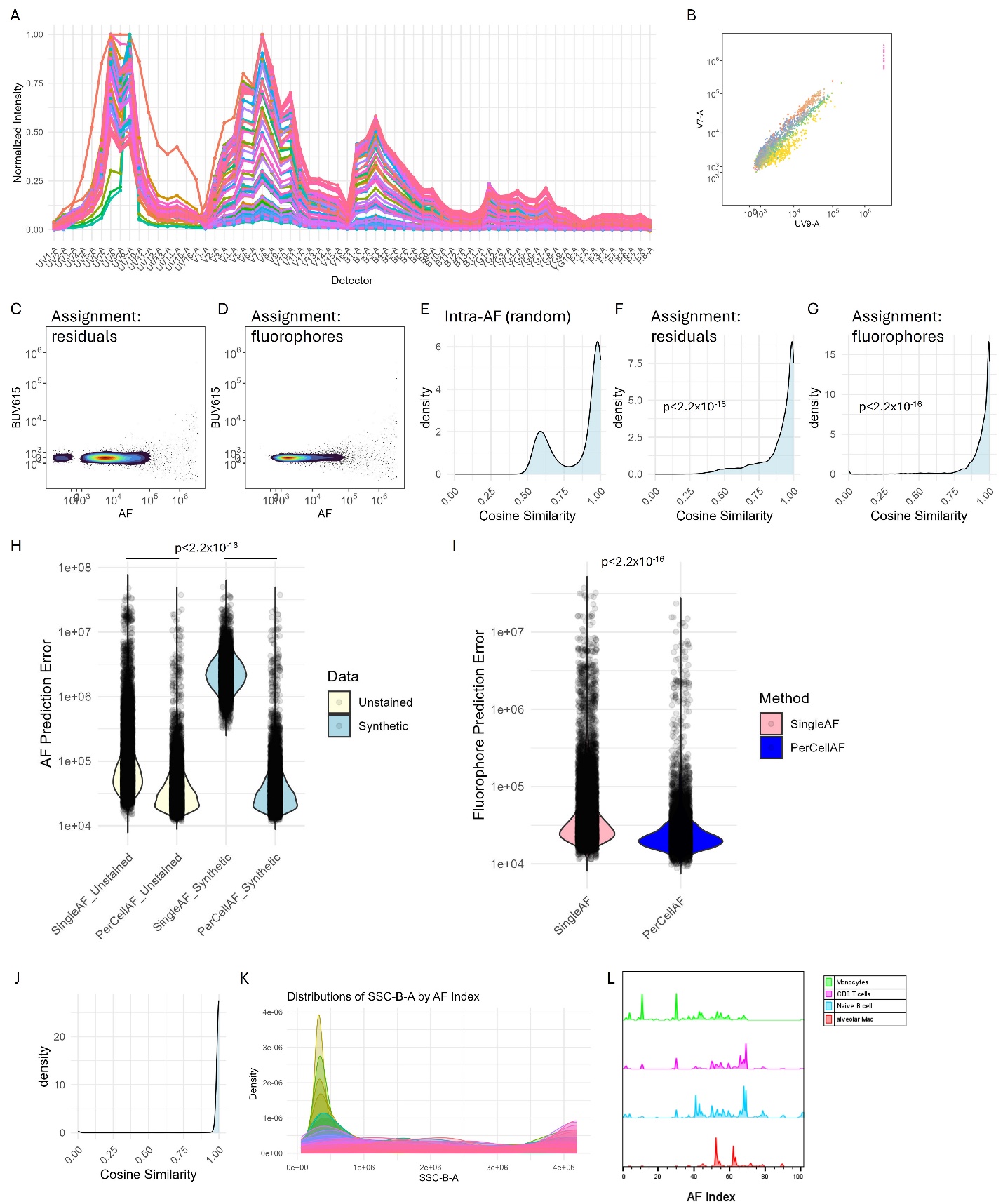

**Supplementary Figure 2. Unmixing of synthetic data with known ground truth using per-cell fluorophore optimization.** Synthetic data were generated in silico by combining data from single-stained cells. In this 42-colour data set, BUV661 and APC are used to detect CD19 and Foxp3, markers that are not co-expressed. We created a set of data *in silico* where the raw data for BUV661^+^ and APC^+^ events (based on OLS unmixing) were combined or maintained as separate original events. In this way, ground truth is available for BUV661^+^, APC^+^ and BUV661^+^APC^+^ events. **A)** Comparison between OLS and per-cell fluorophore optimized unmixing. Frequencies shown are where populations suffer from misclassification with respect to a positivity threshold set based on a value double that of the 99.5^th^ percentile on the unstained sample. **B)** Distribution of cosine similarity values between the model’s predictions for BUV661 and the actual data for BUV661 in cells known to be BUV661+. A value of 1 indicates identity (perfect match).

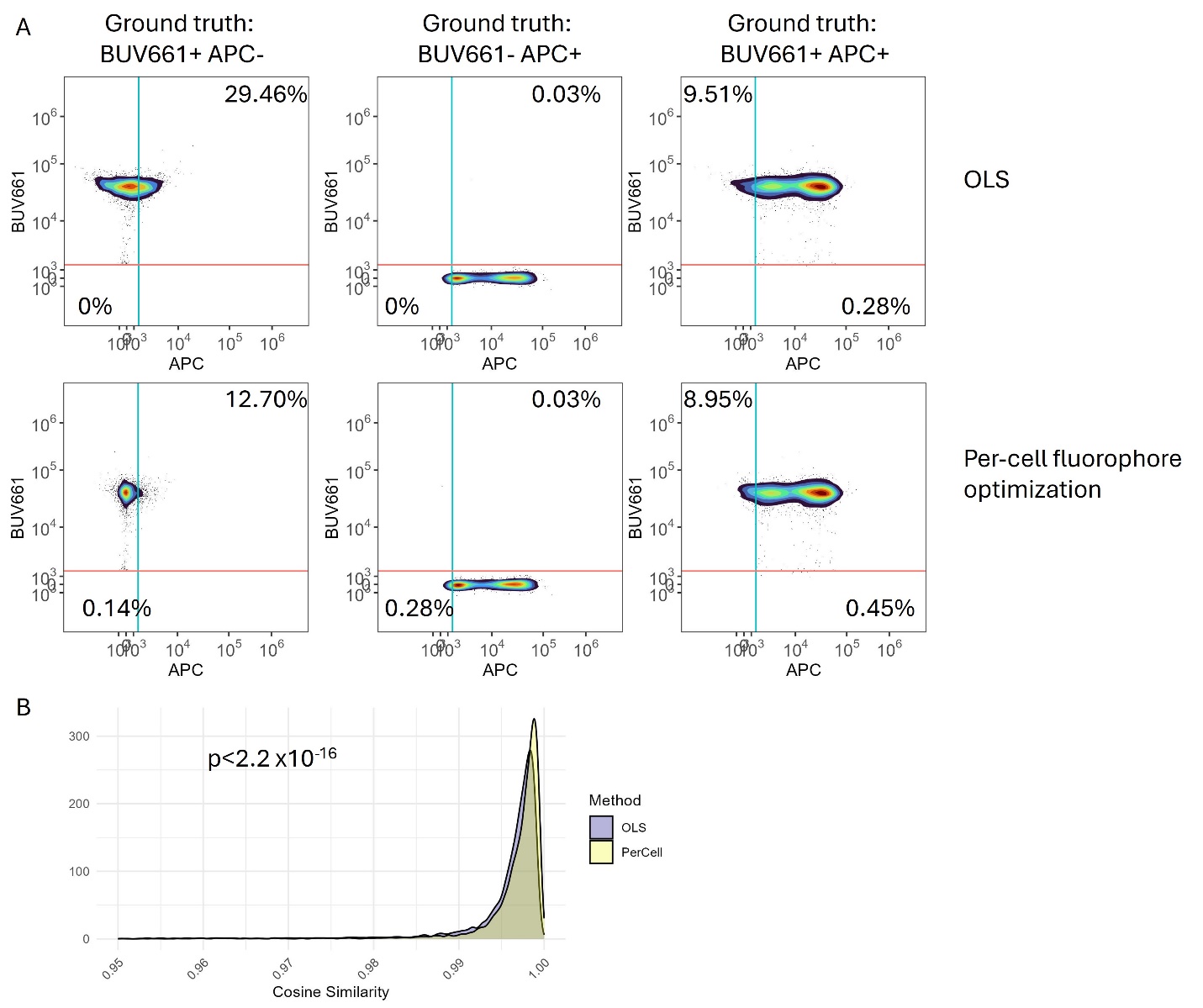
